# A Thermodynamic Limit on the Role of Self-Propulsion in Enhanced Enzyme Diffusion

**DOI:** 10.1101/451682

**Authors:** Mudong Feng, Michael K. Gilson

## Abstract

A number of enzymes reportedly exhibit enhanced diffusion in the presence of their substrates, with a Michaelis-Menten-like concentration dependence. Although no definite explanation of this phenomenon has emerged, a physical picture of enzyme self-propulsion using energy from the catalyzed reaction has been widely considered. Here, we present a kinematic and thermodynamic analysis of enzyme self-propulsion that is independent of any specific propulsion mechanism. Using this theory, along with biophysical data compiled for all enzymes so far shown to undergo enhanced diffusion, we show that the propulsion speed required to generate experimental levels of enhanced diffusion exceeds the speeds of well-known active biomolecules, such as myosin, by several orders of magnitude. Furthermore, the minimum power dissipation required to account for enzyme enhanced diffusion by self-propulsion markedly exceeds the chemical power available from enzyme-catalyzed reactions. Alternative explanations for the observation of enhanced enzyme diffusion therefore merit stronger consideration.

## Introduction

The apparent diffusion coefficients of various enzymes, as measured typically by fluorescence correlation spectroscopy, have been observed to increase in the presence of substrate by as much as 15% to 80%, depending on the enzyme, at maximal substrate concentration. Examples include F0F1-ATP synthase [1], T7 RNA polymerase [2], T4 DNA polymerase [3], bovine catalase [4, 5], jack bean urease [6, 4, 5], hexokinase [7], fructose biophosphatase aldolase [8, 7], alkaline phosphatase [5] and acetylcholinesterase [9]. However, the mechanisms underlying these observations remain largely unexplained. For some enzymes, further experimentation has ruled out certain potential mechanisms for this phenomenon of enhanced enzyme diffusion (EED), including one mediated by local pH changes [6], and propulsion by bubble formation [4]. In a number of cases, the increase in diffusion coefficient relative to baseline has been found to be approximately proportional to the catalytic rate of the enzyme, with a Michaelis-Menten relationship to substrate concentration [5]. This proportionality has naturally led to the suggestion that the chemical reaction catalyzed by the enzyme is a driver of the diffusion enhancement. Indeed, larger, synthetic Janus particles are propelled by the catalysis of reactions at one face of the particle and not the other [10]. Accordingly, a number of possible mechanisms for catalysis-driven self-propulsion of enzymes – i.e. for the transduction of the reaction free energy into mechanical propulsion – have been proposed. These include mechanical swimming [11, 4], pressure waves generated by exothermic reactions [5], and self-diffusiophoresis [12]. However, these specific mechanisms of EED have been debated [13, 8], and none have been proven. Here, we step back from specific propulsion mechanisms and instead analyze the kinematics and thermodynamics of enzyme self-propulsion generically.

## Methods

The degree to which translational diffusion is enhanced may be expressed as:

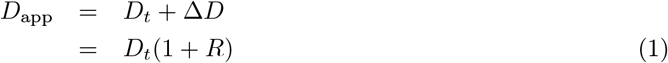

where Δ*D* = *D_app_* – *D_t_* is the difference between the observed, or apparent, diffusion constant, *D_app_*, and the baseline diffusion constant in the absence of enhancement, *D*_t_. Thus, *R* is the relative diffusion enhancement. We consider an enzyme that, within each catalytic cycle, self-propels for a time *t_p_* ≤ *t_c_*, where *t_c_* is the enzymologic turnover time and reciprocal of turnover rate. The magnitude of the propulsive force, *F*, is considered to be constant during *t_p_*. (The consequences of a more complex time-dependence are considered in the Appendix.) For an enzyme in liquid water, the Reynold’s number is very low. Therefore, the dynamics of the enzyme are overdamped, and the propulsion velocity has a constant magnitude *ν* ∝ *F* while the propulsion is active. The vector of the propulsive force and velocity is considered fixed within the enzyme’s internal frame of reference, but it reorients continuously in the lab frame due to rotational Brownian motion of the enzyme. The enzyme is modeled as a hard sphere with radius *a*, moving in liquid water with viscosity *η*, so that the Stokes-Einstein equations may be used to estimate *D_t_* and the rotational diffusion coefficient *D_r_*:

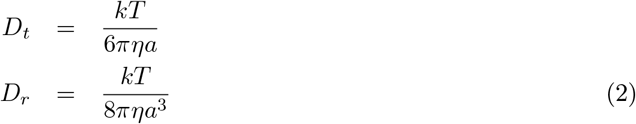

Analytical solutions of the overdamped Langevin equation for self-propelled particles have been developed by Hagen et al. [14], under the assumption that the Stokes-Einstein equations hold and that rotational and diffusional translation are not coupled to each other. In EED experiments, the diffusion coefficient is measured over times much greater than the enzyme’s turnover time, which is in turn usually much greater than the rotational relaxation time of the enzyme, *τ* = (2*D_r_*)^−1^ ∈ [10^−9^s, 10^−6^s] [15]. In this setting Hagen et al.’s Eq 34 applies and yields the mean square displacement as a function of time:

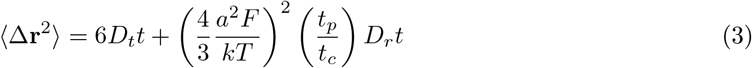

where the first term give the mean-square displacement in the absence of propulsion and the second term captures the effect of propulsion. We have inserted the term *t_p_*/*t_c_* to account for the fact that self-propulsion acts to raise the diffusion constant only during this fraction of the time (see Appendix). Recognizing that *D_app_* = 〈Δr^2^〉/ (6*t*), using Eq 2, and employing Stokes law, *F* = 6*πηaν* to replace force with velocity, one may rewrite Eq 3 as

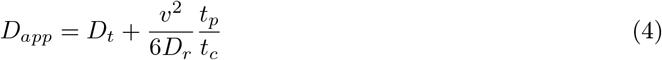

The first term is the contribution of normal Brownian motion, and the second term is the contribution from self-propulsion. The enhancement ratio, *R*, then is

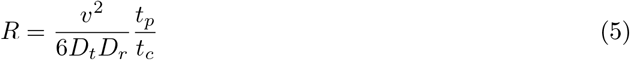

Thus, the propulsion speed required to achieve a given level of diffusion enhancement *R* is given by

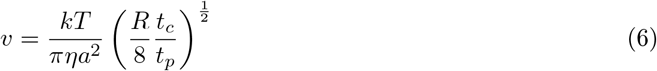

To determine the power required for a self-propelled particle to achieve observed levels of enhanced diffusion, it is necessary to address the energetic efficiency of the self-propulsion mechanism. Rather than make any mechanistic assumptions here, we make the most conservative assumption –i.e., the one requiring least power – by using the minimum energy dissipation theorem. This says that, at low Reynolds number, no propulsion mechanism is more efficient than dragging the particle by external force in a Stokes flow [16, 17, 18, 19, 20]. Accordingly, we consider the power to drag an enzyme molecule in a Stokes flow at the propulsion speed required to generate enhanced diffusion with a specific value of *R*. Inserting *ν* from Eq 6 into Stokes law, *F* = 6*πηaν*, we obtain the required power averaged over the full catalytic cycle:

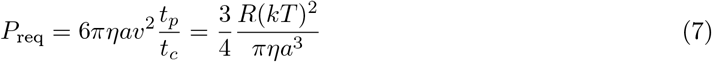

## RESULTS

We first apply Equation 6 to estimate the propulsion speeds needed to account for experimentally observed diffusion enhancements. The minimum thrust speed, *ν*_min_, that would explain the diffusion enhancement is obtained by setting *t_p_* = *t_c_*, because larger speeds are required when *t_p_* < *t_c_*. Given T=298K, the viscosity of liquid water, and a typical enzyme diffusion enhancement of *R* = 0.2 [5], one obtains *ν*_min_(m/s) = 0.21*a*^−2^ (*a* in nm). This quantity depends only on the radius of the enzyme. For catalase, *a* = 5.3nm [21], so the minimum propulsion speed *ν*_min_ = 7 × 10^−3^m/s. Similar values of *ν*_min_ are obtained for the other enzymes that showed EED in experiments, because their radii are similar to catalase (Table 1). These speeds, which amount to ~ 10^6^ enzyme radii per second, are strikingly high. Furthermore, we anticipate that any thrust generated by enzymatic catalysis will persist only for a small fraction of the enzymologic turnover time; i.e., in all likelihood *t_p_* ≪ *t_c_*. As a consequence, based on Eq 6, even higher propulsion speeds would be needed during the short *t_p_* intervals to explain observed values of *R*.

**Table 1:** Turnover rates, experimental hydrodynamic radii, minimum thrust speeds, and required reaction free energies, of enzymes reported to show EED. For multimeric enzymes turnover rate is of the whole multimer. Citations are parenthesized.

| Enzyme | Turnover rate/s<sup>-1</sup> | Radius/nm | $v_{\min}/\text{m s}^{-1}$ <sup>a</sup> | $-\Delta G_{\text{req}}^{\circ}/\text{kJ mol}^{-1}$ <sup>b</sup> |
| --- | --- | --- | --- | --- |
| T4 DNA polymerase | 0.5[22]<sup>d</sup> | 4.6[23] | $1 \times 10^{-2}$ | $1 \times 10^7$ |
| Aldolase | 5[8]<sup>c</sup> | 4.9[21] | $9 \times 10^{-3}$ | $8 \times 10^5$ |
| T7 RNA polymerase | 4[24]<sup>d</sup> | 8.4[25] | $3 \times 10^{-3}$ | $2 \times 10^5$ |
| Hexokinase | 300[7]<sup>c</sup> | 6.3[26] | $5 \times 10^{-3}$ | 6000 |
| ATP synthase | 1000[27]<sup>d</sup> | 6.6[1] | $5 \times 10^{-3}$ | 2000 |
| Alkaline phosphatase | 3000[5]<sup>c</sup> | 7.7[28] | $3 \times 10^{-3}$ | 400 |
| Catalase | 10000[5]<sup>c</sup> | 5.3[21] | $7 \times 10^{-3}$ | 300 |
| Urease | 10000[5]<sup>c</sup> | 7.0[29] | $4 \times 10^{-3}$ | 100 |
| Acetylcholinesterase | 20000[30]<sup>d</sup> | 8.8[31] | $3 \times 10^{-3}$ | 40 |

<sup>a</sup> For $R = 20\%$ diffusion enhancement, using Eq.6 with $t_c = t_p$ .
<sup>b</sup> For $R = 20\%$ diffusion enhancement, using Eq.7 divided by turnover rate.
<sup>c</sup> Turnover rate at $R = 20\%$ , read from the corresponding publication of EED measurements.
<sup>d</sup> Turnover rates of these enzymes were not reported in the publications of their EED measurements, so we instead use the turnover rate when substrate concentration equals $K_m$ , i.e., monomer $k_{cat}$ times number of catalytic sites times 0.5.

Although implausibly high propulsion speeds would be needed to account for EED by self-propulsion, this analysis remains consistent with the observation that larger particles, e.g. Janus particles, can achieve substantial enhancements of diffusion via self-propulsion [10]. This is because, for larger particles, a given propulsion velocity leads to higher values of *R*, mainly through the dependence of *D_r_* on size. Intuitively, the longer the rotational correlation time, the greater the effect of propulsion on the root mean square displacement. Thus, self-propulsion is much more effective at enhancing the diffusion of large particles than that of small particles, such as enzymes.

We now turn to the power required to explain EED and its relation to the chemical energy available from catalysis, which is approximated by the standard free energy of reaction, as explained in the third part of the Appendix. For catalase, with *a* = 5.3nm, the result is *P*_req_ = 3 × 10^6^ kJs^−1^ mol^−1^. The turnover rate of catalase is about 10^4^s^−1^ under conditions which yield a *R* = 20% [5], so this power requirement corresponds to a minimum required reaction free energy of 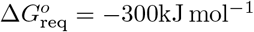. This is well above the standard free energy of reaction, Δ*G*° = −95kJ mol^−1^, computed from the standard free energies of formation of the reactant and products [32]. For enzymes with lower turnover rates, the required reaction free energies range up to 5 × 10^6^kJmol^−1^ (Table 1). These required reaction free energies are far larger than what is available from the free energy of the chemical reactions catalyzed by the enzymes. For example, for alkaline phosphatase, Δ*G*° = −8.5kJmol^−1^; for urease, Δ*G*° = −20kJmol^−1^; for acetylcholinesterase, Δ*G*° = −17kJmol^−1^[33]. The magnitudes of the reaction free energies in Table 1 may be put into perspective by considering that the standard free energy of hydrolysis of ATP, the cell’s energy currency, is only about −32 kJ mol^−1^ [34]. Furthermore, as detailed in the Discussion, the power requirements derived here are conservative, and the actual power requirements probably exceed what is available by an even larger margin. Thus, it is unlikely that experimental observations of enhanced enzyme diffusion can be accounted for by catalysis-driven self-propulsion.

## Discussion

We now critically examine the approximations and assumptions used in the present theory and consider the results in light of recent relevant experimental studies.

Three key assumptions in this analysis are conservative, in the sense of lowering the estimate of the power required to achieve a certain level of enhanced diffusion. First, we used the minimum energy dissipation theorem, based on the assumption of a Stokes flow around the enzyme, to estimate the minimum power required for a given propulsion velocity. Any real propulsion likely generates a non-Stokes flow field around the enzyme, resulting in higher viscous dissipation integrated over whole space than in the ideal Stokes flow, and hence to lower efficiency than assumed here. (Intuitively, if one replaces the enzyme by a bacterium, we computed the dissipation associated with pulling it through the water with an optical trap, rather than the greater dissipation associated with its using flagellae to swim at the same speed.) Indeed, a bacterial propulsion mechanism, which, unlike a non-motor enzyme, has been optimized during evolution, was found to have only about 1% of the maximum propulsion efficiency associated with pure Stokes drag [18]. Additionally, a propulsion mechanism might rely on local chemical gradients, imposing an additional entropy production term as the chemical gradients spontaneously dissipate. Thus, although we have used the maximum efficiency assumption, the true efficiency of any enzyme propulsion mechanism is probably orders of magnitude lower. This makes it even less probable that the required power could be provided by the available chemical energy.

Second, we assumed that the propulsion mechanism increases the apparent translational diffusion coefficient without increasing the enzyme’s rotational diffusion coefficient, *D_r_*. We are not aware of any experiments that report on the rotational diffusion rates of enzymes undergoing translational EED, but any translational propulsion mechanism would probably also increase the rate of rotational diffusion. This is because there is no reason to expect that a propulsive force will not also exert a torque and thus drive rotation. In fact, the rotational diffusion coefficient of 30nm Pt-Au Janus particles increases by up to 70% when they are catalytically active and undergoing enhanced translational diffusion [10]. This is relevant here because, as evident from Eq 4, increasing *D_r_* would further increase the velocity *ν* needed to achieve a given level of *D_app_*. Therefore, even more power would be required, again making EED harder to explain on the basis of catalysis-driven self-propulsion. In addition, any chemical energy expended in driving rotational motion would become unavailable to drive translational motion.

Third, we assumed that the thrust speed *ν* is constant during the interval *t_p_*. Using Fourier analysis, we show in the Appendix that allowing *ν* to be time-varying during *t_p_* could only increase the thrust speed and power required to achieve a given degree of diffusion enhancement. The Appendix furthermore proves a more implicit but intuitive assumption we have used, that diffusion enhancement and power requirements scale linearly with the duty ratio *t_p_*/*t_c_*. This is found to hold as long as any significant high frequency components in the thrust velocity are slow relative to the rotational diffusion time of the enyzme. Intuitively, if a high frequency component of the thrust speed reverses direction before the enzyme has had time to reorient, the motion due to this component can be canceled in the lab frame, leading to a minimal contribution to the net translational displacement. In contrast, if the enzyme has time to rotate before the thrust component reverses, the reversed component will act in a different direction in the lab frame, leading to less cancellation and more net displacement.

Several other assumptions also deserve comment. Our use of the Stokes-Einstein equations with stick boundary conditions is justified by several considerations. First, changing to slip boundary conditions would merely replace the factor of 1/6 in the Stokes-Einstein equation by a factor of 1/4, which would not change our conclusions. Additionally, simulations of spherical macromolecule-sized particles in solution yield translational diffusion coefficients that are bracketed by the results of the Stokes-Einstein equation computed with stick and slip boundary conditions, using the geometric radii of gyration of the solutes [35]. And if one assumes stick boundary conditions in mapping from measured translational diffusion coefficients of proteins in water to effective radii and then from radii to the predicted rotational diffusion coefficient, the results agree with experiment to within about 50% [21]. Interestingly, the actual rotational diffusion constants tend to be higher, rather than lower, than those predicted by Eq 2 [36]. Correcting in this direction would only strengthen our conclusions, because increasing *D_r_* means that even more power is required for a given value of *R*. Finally, treating the enzymes for which enhanced diffusion has been observed as spherical is reasonable for these globular proteins; highly nonspherical, e.g. rod-like, proteins, may deserve further analysis.

Additionally, we have treated each enzyme molecule’s motion as independent of the motions of the other enzymes in solution. We tested this assumption by applying the hydrodynamic interaction model of Mikhailov et al. [37] to the case of enzymes at the very low concentrations, about 10 nM, used in typical EED measurements. The resulting hydrodynamic interactions are found to be negligibly small.

It is worth considering the present analysis in the context of recent, high-resolution experimental studies of EED. In two elegant studies, Jee et al. combined stimulated emission depletion microscopy with fluorescence correlation spectroscopy to study enzyme diffusion at very high spatial resolution [9, 38]. Intriguingly, when urease in the presence of urea was studied with a small beam waist (50-250 nm), a fast component of translational motion was revealed. The authors interpreted the fast component as being the result of propulsive motion powered by the urease reaction and argued that this self-propulsion could explain enhanced diffusion of urease. Perhaps the chief reason for the difference in their conclusion relative to ours is that their rotational diffusion time of 2.9-5.6*μ*s corresponds to a hydrodynamic radius *a*=10-12nm, which is considerably larger than the value of 7.0nm reported in a prior experimental study [29] and used here. The smaller hydrodynamic radius used here is further supported by our analysis of the hexameric biological unit of urease [39] with the program HYDROPRO [40], which yields translational and rotational diffusion coefficients corresponding to hydrodynamic radii of 6.6nm and 6.7nm, respectively. In addition, one may infer the hydrodynamic radius of urease from the baseline translational diffusion coefficient of 29*μ*m^2^/s reported by Jee et al; the result is 7.5nm, which is close to the value we used. Based on Eq 7, going from a radius of 10-12 nm to the more plausible value of 7nm used here leads to a three to five fold increase in the power requirement for a given degree of diffusion enhancement *R*. Given that Jee and coworkers’ estimated value of the free energy required for each catalytic cycle, 25kJmol^−1^ [38], is already slightly higher than the reaction free energy of this enzyme, 20kJ mol^−1^ [33], an upward adjustment based on this consideration makes it difficult to support the hypothesis that catalytic self-propulsion explains EED in urease. Interestingly, their reported rotational diffusion time of 44-46*μ*s for the enzyme acetylcholinesterase corresponds to a hydrodynamic radius of about 25nm, which is about three times the experimentally determined hydrodynamic radius of the largest globular form of this enzyme [31, 41]. One may speculate that the abnormally low rotational reorientation rates inferred by Jee et al. could reflect extrinsic perturbations of the enzymes, such as fluid flows, varying on a timescale of about 10*μ*s.

## CONCLUSIONS

The present analysis shows that the propulsion speeds required to explain experimentally observed levels of EED by the mechanism of catalytic self-propulsion are implausibly large. More fundamentally, the power levels needed to to account for observed levels of diffusion enhancement by catalytic self-propulsion are greater than those available from enzyme-catalyzed chemical reactions. For most enzymes, the power requirement is orders of magnitude too great, and even for the faster enzymes, the power required is still considerably larger than that afforded by the reaction. Moreover, the power actually required to generate observed levels of diffusion enhancement is probably greater than our estimates, because we have used conservative approximations that lead to lower estimates of the required power. However, because the power required for a given level of diffusion enhancement decreases sharply with increasing particle size, our results remain consistent with experimental observations that self-propulsion of micron-scale particles with surfaces coated with a metallic catalyst [10] or with immobilized enzymes [42] leads to significantly enhanced translational diffusion. The propulsion direction of larger particles randomizes more slowly, so the contribution of propulsion to translational diffusion is increased. We conclude that enhanced diffusion of enzymes cannot easily be explained by self-propulsion powered by the chemical energy of the catalyzed reactions.

It is of interest to consider other explanations for EED. One possibility is an increase in normal, thermally driven, translational diffusion. This could result from a decrease of the mean hydrodynamic radius of the enzyme in the course of the catalytic cycle, as recently noted [8, 43]. Alternatively, it has been proposed [44] that the catalytic cycle might raise the temperature of nearby solvent enough to increase the enzyme’s diffusion constant, through *η* and *T* in Eq.2. However, the viability of this explanation appears to rely on use of the thermal conductivity of air, rather than water [44], as the effect becomes negligible when the thermal conductivity of water is used. Global heating of the solution due to release of chemical energy is also insufficient to explain observed diffusion enhancement [5, 45]. It is worth noting, too, that exothermicity, and even chemical catalysis itself, is not required for at least some reported instances of EED [8, 2].

Thus, the mechanisms of EED remain obscure. Further experimental studies may help solve this puzzle. It has been suggested [46, 13] that fluorescence correlation spectroscopy measurements may be subject to experimental artifacts, such as subunit dissociation and fluorophore quenching, so that further controls, such as those employed by Jee and coworkers [9, 38], are of high value. Because the turnover rate is needed to convert the power requirement (Eq. 7) to 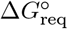 and compare with standard thermodynamic data, it would also be helpful if the turnover rate of the enzymes being studied could be measured under the precise conditions of each diffusion study, to avoid uncertainties that may result from literature data measured under different conditions and from reliance on an assumption of Michaelis-Menten kinetics. Alternative technologies for measuring diffusion enhancement may also provide different perspectives. For example, although FCS studies of aldolase demonstrated EED [7, 8], aldolase did not show enhanced diffusion when studied by dynamic light scattering [47] or by NMR [48]. On the other hand, an electroechemical experiment has providing supporting evidence of catalase EED [49]. Intriguingly, a study in which enzyme molecules were confined to an approximately 2D region to enable single-molecule tracking showed strong enhanced diffusion, though we note that interpretation of these data is complicated by the fact that the baseline diffusion coefficients were markedly reduced relative to their 3D values[50]. Further direct tracking studies [51] could be useful both to confirm the phenomenon of EED and to provide details that might bear on mechanism.

## Appendix

### Fourier analysis of time-varying thrust – general analysis

The derivation in the main text treats the self-propulsion thrust as constant during an interval *t_p_* within each catalytic cycle of duration *t_c_* ≥ *t_p_*. Here, we examine the consequences of a more general time-varying thrust. We make the reasonable assumption that the time over which the translational diffusion constant is measured, *t_m_*, is much larger than the duration of catalytic cycle, *t_c_* (ms to s), which in turn is is much larger than the rotational relaxation time *τ* ≡ (2*D_r_*)^−1^ of the enzyme (ns to *μ*s). We address the effect of time-varying propulsion on translational diffusion by expanding the propulsion speed in a Fourier series, as previously done by Lauga in the context of reciprocal swimming [52], and extend the analysis to determine how time-variation affects the efficiency with which propulsive power generates enhanced diffusion.

Consider an enzyme with a time-dependent, self-propulsion speed *ν*(*t*), whose translational diffusion is evaluated from time *t* = 0 until the end of some experimental time, *t_m_*. As in the main text, the direction of the propulsion is fixed in the enzyme’s frame of reference and therefore reorients in the lab frame of reference due to rotational diffusion of the enzyme. After periodic extension, *ν*(*t*) can be expanded into a Fourier series:

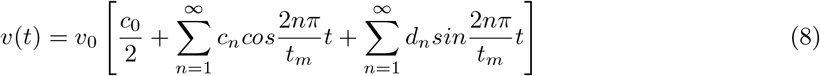

This time-varying propulsion speed generates an increment in the translational diffusion coefficient given by Lauga’s equation 7 [52],

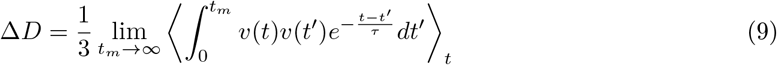

where we have inserted missing angle brackets, indicating an ensemble average over reference time *t* in the integral. This expression yields a well-defined result because *t_m_* ≫ *τ*. Substitution of the Fourier series into this expression yields:

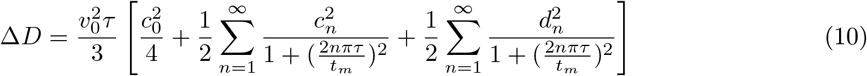

This equation decomposes the diffusion enhancement into contributions from each Fourier component. The mean power consumption, 〈*P*〉 = 6*πηa*〈*ν*(*t*)^2^〉, may similarly be decomposed into contributions from each frequency component,

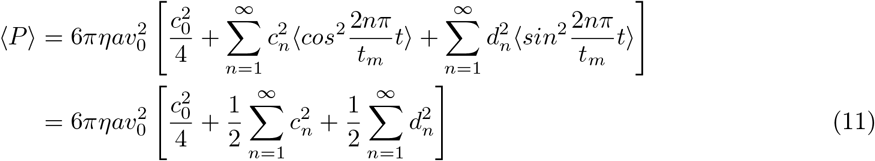

Here, we have used the orthogonality of the Fourier components to eliminate cross terms, and have made the substitutions 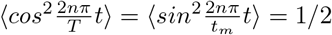.

Comparing Eq.10 and Eq.11 reveals that, given a set of amplitudes *c*_0_, *c*_1_, …, *c_n_*, *d*_1_, *d*_2_, … *d_n_*, higher frequency components (i.e., ones with larger subscripts) generate smaller contributions to the diffusion coefficient but equal contributions to the power consumption. The efficiency of diffusion enhancement, normalized to that for constant propulsion, is given by Eq 10 and Eq 11 as 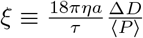. It is apparent from the present analysis that the efficiency is greatest when only the constant thrust component, *c*_0_, is nonzero; i.e., when the thrust speed is constant during the enyzme’s catalytic cycle, as assumed when considering the minimum thrust speed in the main text. Any variation in thrust over time can only reduce *ξ* to below one. Thus, “scheduling” the thrust can not decrease the power needed for a given level of diffusion enhancement to below the power needed for constant thrust.

### Fourier analysis of time-varying thrust – square-wave case

In the main text, we assumed a square wave thrust schedule, with constant nonzero thrust during *t_p_* < *t_c_* and zero thrust during the rest of *t_c_*. We argued that the diffusion enhancement and the minimal power dissipation both scale linearly with the duty ratio *t_p_*/*t_c_*. For diffusion enhancement, it should be apparent that this holds, because the ensemble average in Eq 9 is proportional to the portion of time when *ν*(*t*) is non-zero. Nonetheless, it is of interest to confirm these arguments numerically within the Fourier analysis. To do this, we consider the speed to be *ν*(*t*) = *ν*_0_ when 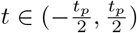, and *ν*(*t*) = 0 elsewhere in 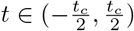. The corresponding Fourier series is:

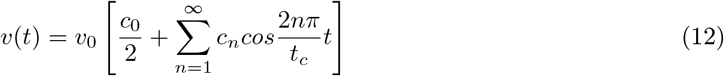

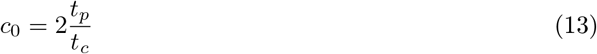

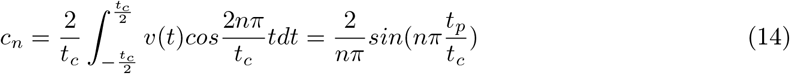

Inserting these expressions into Eq 10, with *τ*/*t_c_* = 0.01, which corresponds to the case of urease, yields the expected linear variation of Δ*D* with *t_p_*, as shown in Figure 1.

**Figure 1:**
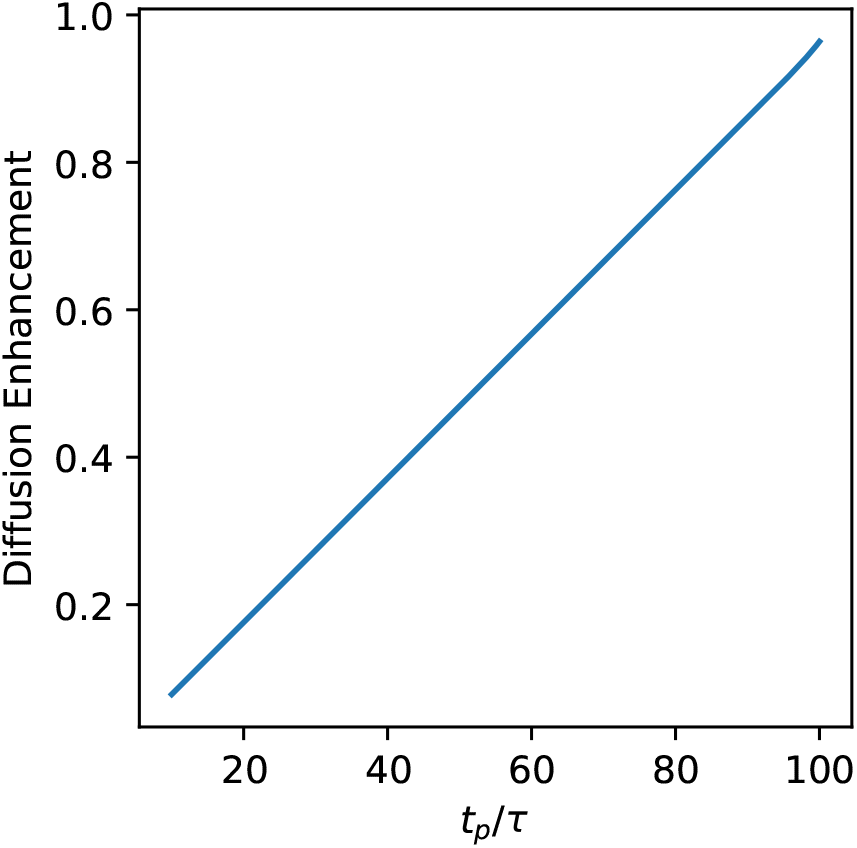
Numerical evaluation of the diffusion enhancement given by Eq 10 confirms the linear relationship between the ratio *t_p_*/*τ* and the diffusion enhancement for a square-wave thrust schedule. The enhancement is plotted relative to the case *t_p_* = *t_c_*. The value of *t_c_* and *τ* correspond to urease from Table 1.

We next examine the efficiency, ξ, for this square wave thrust schedule:

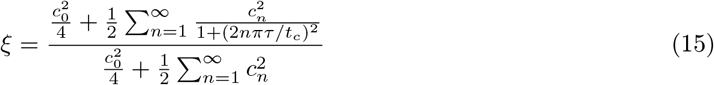

For constant speed with *t_p_* = *t_c_*, this yields *ξ* = 1. The loss in efficiency when *t_p_* < *t_c_* then is given by:

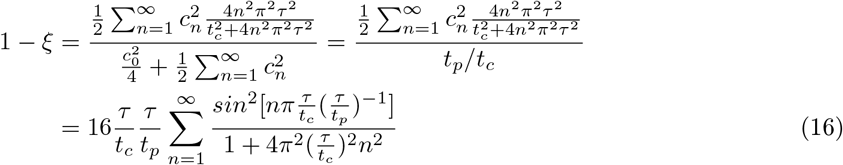

where we have used Parseval’s theorem to evaluate the denominator and then inserted Eq 14. For given values of *τ* and *t_c_*, the maximum drop in efficiency is expected to happen when *t_p_* is much smaller than *t_c_*, as this increases the weight of the high frequency components of the thrust velocity. Focusing, then, on this low-efficiency limit, we can approximate the summation with an integral and then evaluate the integral using the residual theorem:

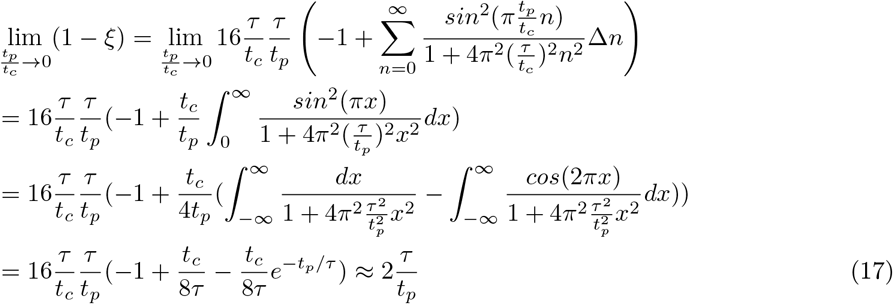

Consequently, as *t_p_* ≫ *τ* is expected for most enzymes, the efficiency will remain near unity, even under the extreme assumption that *t_p_* ≪ *t_c_*. This result supports our approximation in the main text that the diffusion enhancement caused by a square-wave thrust schedule is proportional to *t_p_*. It also shows that our assumptions are conservative, because not invoking this approximation would decrease the efficiency and further increase the power requirement. The analytical result in Eq 16 is elaborated by numerical calculations of the drop in efficiency ξ, as drawn in Figure 2. Here, *t_c_*/*τ* spans the range of this ratio found for the enzymes in Table 1, from 100 for urease to 3 × 10^6^ for DNA polymerase. Three values for *t_p_*/*τ* are used, subject to the requirement that *t_p_* < *t_c_*.

**Figure 2:**
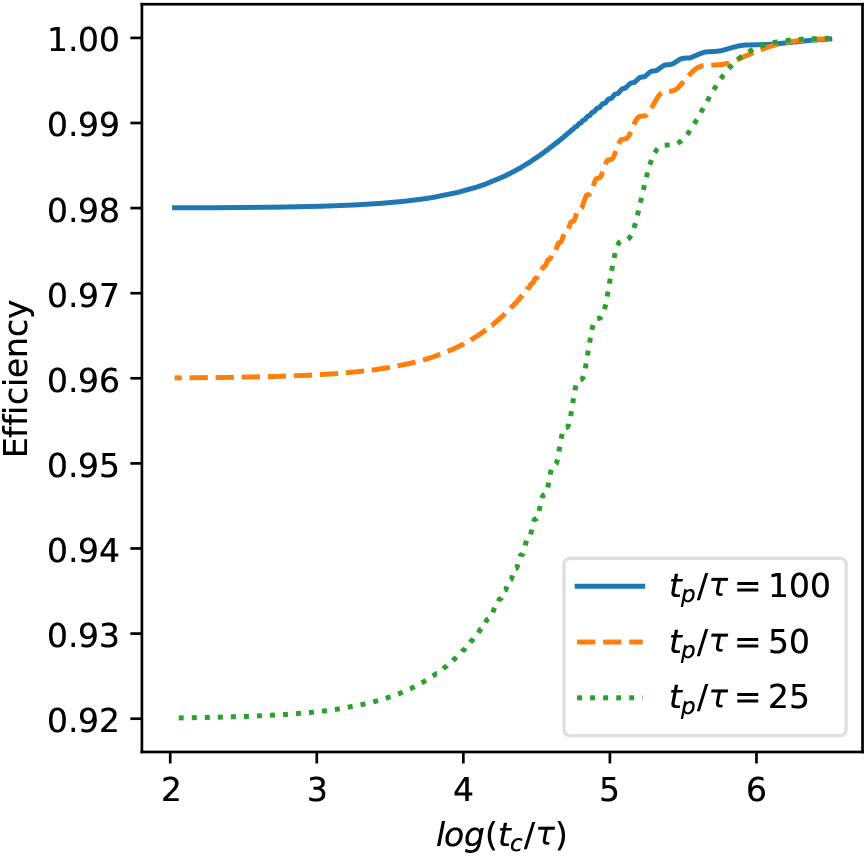
Relationship between efficiency *ξ*, and the ratio *t_c_*/*τ*, plotted for three values of *t_p_*/*τ*.

The near proportionality of both Δ*D* and 〈*P*〉 to *t_p_* may be understood more intuitively by reference to Eq 10 and Eq 11. Because the denominator in Eq 10, 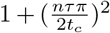, is near unity except for very large *n*, low-frequency components deviate only very slightly from the zeroth component in efficiency. On the other hand, high-frequency components with large *n* have negligible amplitudes, as 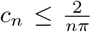, so they do not alter efficiency either. Therefore, the diffusion enhancement and the power requirement both scale near-linearly with duty ratio *t_p_*/*t_c_*, leading to near-uniform efficiency. It is of interest to note, however, that efficiency would fall if there were significant oscillations in *ν*(*t*) on the timescale of *τ* or smaller. In this regime, the non-zero velocity components reverse direction before the enzyme has had time to rotate, so there is little net displacement due to the thrust. In contrast, when the non-zero velocity components do not reverse until the enzyme has had time to rotate, the net effect of the time-varying thrust is to generate randomly directed displacements, which contribute to the apparent diffusion constant.

### Free energy for self-propulsion available from an enzyme-catalyzed chemical reaction

In the main text, we took the standard free energy of the reaction, Δ*G*°, to be the free energy from an enzyme-catalyzed chemical reaction that is available to power the enzyme’s self-propulsion. A concern with this approach may be that, when the two sides of the chemical reaction have different numbers of solute molecules, Δ*G*° depends on the arbitrary standard concentration, *C*°, and the available free energy ought not depend on an arbitrary quantity. Here, we show that the standard reaction free energy is, in fact, a good approximation to the free energy available from the combined processes of substrate-enzyme binding, chemical reaction, and product release, so long as the standard concentration is set to its customary value of 1 mol/L. This section thus justifies the use of the standard concentration in the main text, while also offering insight into how more refined estimates of the available free energy might be made.

First, it is instructive to consider whether it would be appropriate to take the free energy available for propulsion to be the free energy of reaction under the experimental conditions at which enzyme diffusion was studied; i.e., Δ*G* = Δ*G*° + *RT* ln *Q*, where *Q* is the experimental concentration quotient, assuming activity coefficients near unity. This approach is problematic, because it would require a physical mechanism that could couple the macroscopic concentrations of substrate and product to the the local events at a single enzyme molecule. Instead, if one considers the entire catalytic process, from enzyme-substrate encounter through release of product to the bulk, the only steps that could contribute free energy to enzyme propulsion are those in which the enzyme interacts significantly with the substrate or product. Such interactions occur only when the substrate or product molecules are near the enzyme, so the free energy available for propulsion may be termed the local free energy, Δ*G*_loc_.

The local free energy may be estimated with the thermodynamic cycle shown in Fig.3, which illustrates a case where one substrate molecule, S, present at concentration *C_S_*, is converted to two product molecules, P1 and P2, present at concentrations *C*_*P*1_ and *C*_*P*2_, respectively, with a free energy of reaction under experimental conditions of 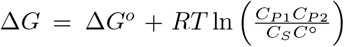, assuming ideal solutions. The lower route of the cycle breaks the process into five steps. In the first, step, the substrate molecule is, in effect, compressed into the region near the enzyme where enzyme-substrate interactions are non-negligible, under the artificial assumption that only steric interactions exist between the two molecules. The volume of this region is termed *V*_loc,S_, and the free energy change associated with this step is Δ*G*_1_ = −*RT* ln (*V*_loc,S_*C_S_*). The subsequent three steps are those for which the free energy change, Δ*G*_loc_, could contribute free energy to propulsion. Here, the non-steric enzyme-substrate interactions are turned on, the substrate is converted to product, and then all non-steric interactions between the enzyme and products are artificially turned off while the products are constrained to remain in the region where these interactions were non-negligible. For products P1 and P2, the volumes of these local regions are, respectively, *V*_loc,P1_ and *V*_loc,P2_. Finally, the constrained products are released to their solute concentrations, with free energy change Δ*G*_2_ = *RT* ln (*V*_loc,P1_*V*_loc,P2_*C*_*P*1_*C*_*P*2_). Closing the thermodynamic cycle now allows one to show that 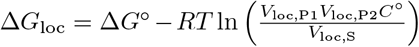. Note that if *V*_loc_ is given in units of nm^3^, then the 1 mol/L standard concentration should be written as 0.6 molecules/nm^3^. The steps corresponding to the local free energy have the character of a unimolecular process, and this quantity is, accordingly, independent of the standard concentration, C°, because any change in C^*∘*^ causes equal and opposite changes in Δ*G*° and the second term of Δ*G*_loc_. If the enzyme interacts with substrate and product molecules over similar ranges, we may write all three local volumes as the same quantity *V*_loc_, and the local free energy takes the simpler form Δ*G*_loc_ = Δ*G*° − *RT* ln(*V*_loc_C°). A straightforward generalization to other stoichiometries yields Δ*G*_loc_ = Δ*G*° − (*N_P_* − *N_S_*) *RT* ln (*V*_loc_*C*°), where *N_P_* and *N_S_* are the numbers of product and substrate solutes, respectively. Again, although Δ*G*° depends on the standard concentration, this dependency is cancelled by the factors of *C*° in the added term.

**Figure 3:**
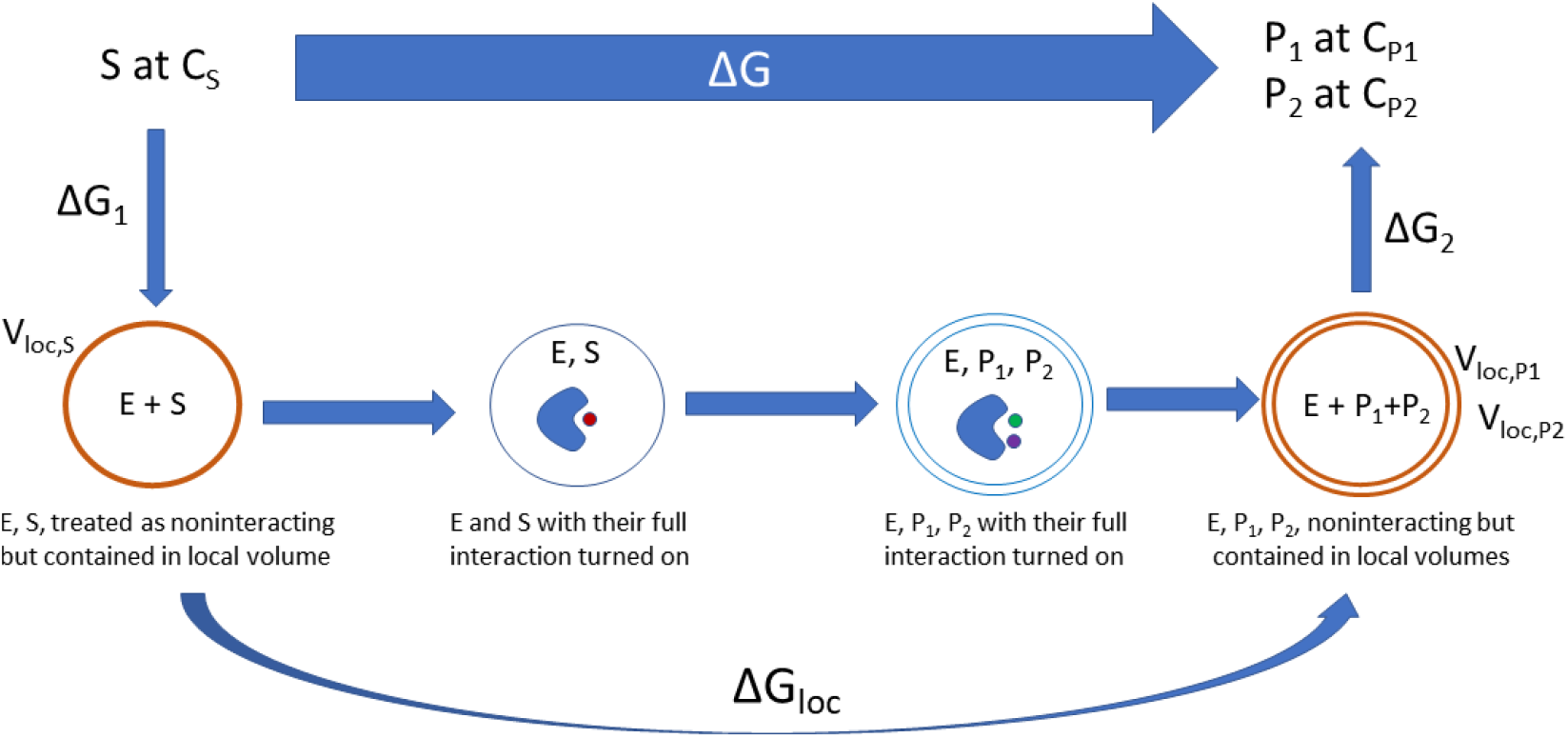
Definition of Δ*G*_loc_ via a schematized thermodynamic cycle. See text for details.

The quantity *V*_loc_ is the volume covered by the interaction range of substrate and product molecules with the enzyme. We estimate this quantity by considering the interaction region to be a hemisphere around the enzyme active site with a 1 nm radius typical of protein-ligand interaction ranges, as determined from molecular dynamics simulations [53, 54]. With these assumptions, *RTln*(*V*_loc_*C*°) = 0.6kJ mol^−1^. Note that this quantity is rather insensitive to the precise choice of V_loc_ because of the logarithm. For urease, where one molecule of urea is decomposed into one carbon dioxide and two ammonia molecules, *N_P_* − *N_S_* = 2, so Δ*G*_loc_ = Δ*G*° − 2*RTln*(*V*_loc_C°) = −20kJ mol^−1^ − 1.2kJ mol^−1^ = −21.2kJ mol^−1^. For acetylcholinesterase, where one molecule of acetylcholine is decomposed into one acetic acid and one choline, *N_P_* − *N_S_* = 1, so Δ*G*_loc_ = Δ*G*° − *RTln*(*V*_loc_*C*°) = −17kJmol^−1^ − 0.6kJ mol^−1^ = −17.6kJ mol^−1^. Thus, the local free energies available to drive propulsion remain close to the standard binding free energies appropriate to *C*° = 1mol/L, as was to be demonstrated. We note that this result is serendipitous, as changing to a different standard concentration would not change Δ*G*_loc_, but would change Δ*G*°.

